# Ecological and functional stratification of the stool microbiome predicts response to immune checkpoint inhibitors across cancer types

**DOI:** 10.1101/2025.05.07.652660

**Authors:** Vera A. Orletskaia, Evgenii I. Olekhnovich

## Abstract

Despite the recognized role of the gut microbiome in modulating immune checkpoint inhibitor (ICI) efficacy, the ecological principles governing this relationship remain elusive. Moving beyond cataloging specific bacteria, we investigated whether general ecosystem properties determine clinical outcome. Through genome-resolved metagenomic analysis, we constructed a comprehensive catalog from 951 stool metagenomes and subsequently analyzed a curated subset of 624 samples from 11 multi-cancer cohorts. Our catalog comprises 3,816 non-redundant metagenome-assembled genomes (MAGs) and reveals key ecological determinants of ICI response. We found that clinical benefit is associated with an ecosystem dominated by prevalent, autochthonous taxa. A taxon’s prevalence in the population positively correlated with its association with positive outcome. Functionally, responder-associated microbes were enriched in genomic capacity for complex carbohydrate metabolism (including specialized mucin degradation) and amino acid biosynthesis. In contrast, non-response was characterized by enrichment of low-prevalence, exogenous (oral and food-derived) bacteria and a functional shift toward nucleoside metabolism, indicative of a dysbiotic state focused on replication. A log-ratio biomarker capturing this ecological shift provided generalizable predictive value across independent cohorts (mean AUC = 0.67 ± 0.13). Our results support an ecological interpretation of the “Anna Karenina principle” in microbiomes: response is linked to a stable, functionally coherent microbial community, whereas non-response represents a destabilized state with high individual variability. This reframes the search for biomarkers from individual taxa to the assessment of ecosystem stability and metabolic competence, providing a foundation for microbiome-targeted strategies to improve cancer immunotherapy outcomes.

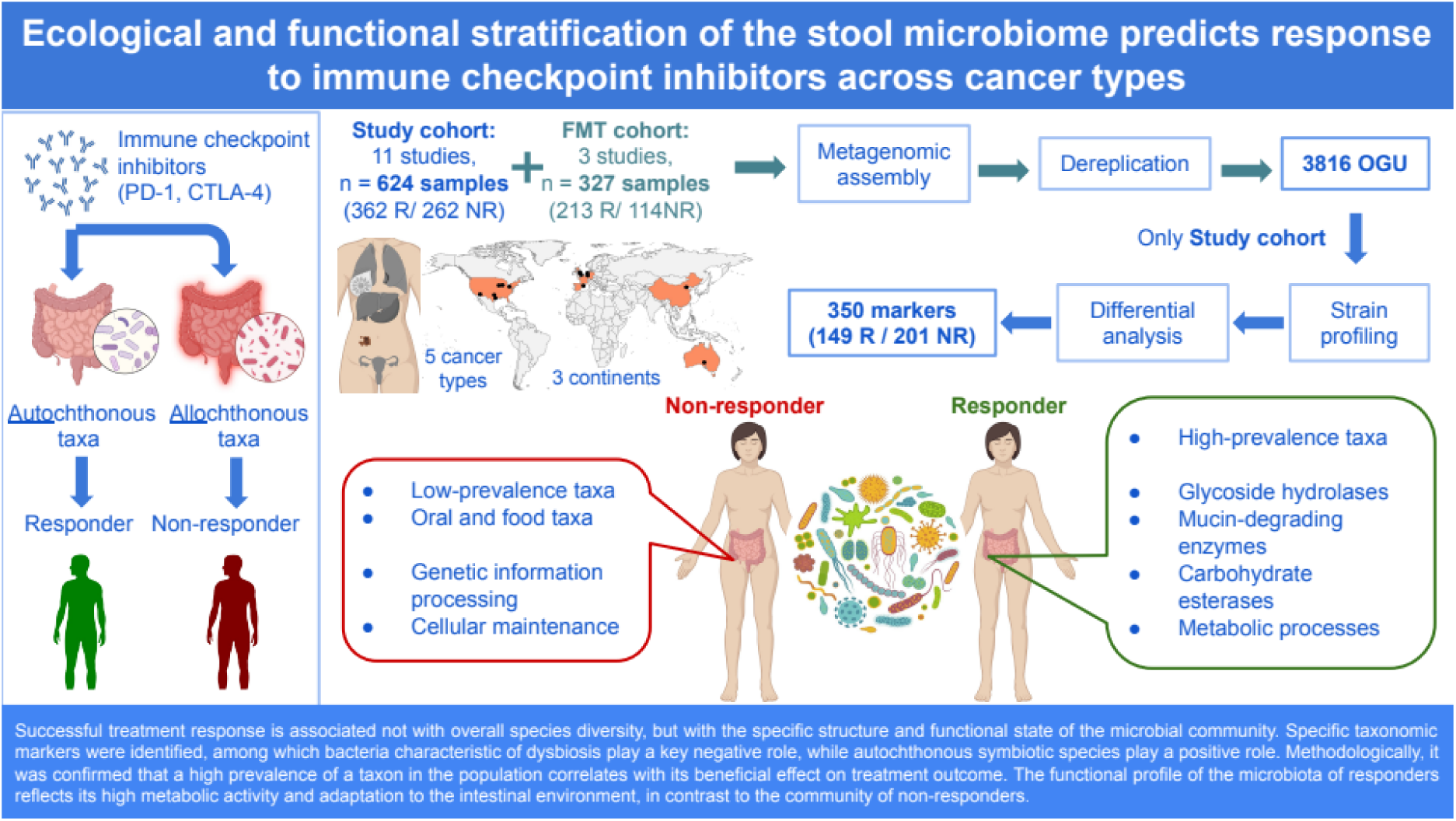

## Introduction

The introduction of immune checkpoint inhibitors (ICIs) targeting cytotoxic T-lymphocyte-associated antigen 4 (CTLA-4) and programmed cell death protein 1 (PD-1) has revolutionized the treatment of advanced malignancies. However, clinical outcomes remain heterogeneous, with a significant proportion of patients exhibiting intrinsic resistance or acquiring resistance after an initial response [Robert et al., 2015]. For instance, ICIs demonstrate limited efficacy in approximately 50% of patients with metastatic melanoma [Larkin et al., 2015] and are associated with a substantial risk of immune-related adverse events [Robert et al., 2019]. This underscores the critical need for predictive biomarkers to optimize patient selection and improve therapeutic outcomes.

Accumulating evidence from preclinical models [Sivan et al., 2015] and clinical cohorts [Routy et al., 2018] indicates a significant association between the composition of the gut microbiome and the efficacy of cancer immunotherapy. The gut microbiome is now recognized as a pivotal modulator of systemic anti-tumor immunity. This link is further supported by interventional studies demonstrating that fecal microbiota transplantation from responding patients can augment ICI response in germ-free mice [Gopalakrishnan et al., 2018] and in human patients [Baruch et al., 2021]. While early research predominantly focused on melanoma [Tsakmaklis et al., 2023], recent efforts are aimed at identifying conserved microbial signatures and mechanisms across diverse cancer types [Gunjur et al., 2024; Lin et al., 2025]. This shift is predicated on the hypothesis that fundamental host-microbiome-immune interactions are broadly relevant to immunotherapy success.

However, despite these advances, the specific biological mechanisms through which the gut microbiota influences immunotherapy response remain incompletely characterized. A recent meta-analysis consolidated evidence for stool metagenomic markers associated with positive outcomes in melanoma immunotherapy [Olekhnovich et al., 2023]. Building on this, our previous application of genome-resolved metagenomics revealed that a reduced functional potential for key microbial metabolic processes - including glycoside hydrolase activity, amino acid metabolism, and acetate synthesis via the Wood-Ljungdahl pathway - was associated with diminished efficacy of immunotherapy in metastatic melanoma [Zakharevich et al., 2024]. However, whether these ecological and functional principles represent generalizable rules across cancer types remains unknown.

By constructing a comprehensive catalog of non-redundant metagenome-assembled genomes (MAGs) and analyzing a large, multi-cancer cohort spanning 624 metagenomic samples from 11 studies across five continents, we established fundamental microbial determinants of immunotherapy response. Through this approach, we discovered that clinical benefit is associated with a conserved ecosystem of prevalent, autochthonous taxa exhibiting genomic capacity for complex carbohydrate metabolism and amino acid biosynthesis. In contrast, non-response manifests as a distinct ecological state characterized by the enrichment of low-prevalence allochthonous microbes (including oral and food-associated bacteria), by the disruption of core biosynthetic pathways, and by a shift toward nucleoside metabolism. This functional profile potentially reflects a move away from host-supportive symbiosis toward a state geared for microbial proliferation. Integration of mixed-effects modeling and machine learning demonstrated that a simple log-ratio-based ecological index derived from these features provides generalizable predictive power (AUC = 0.67 ± 0.13) across diverse cohorts. Collectively, our findings underscore that a stable gut ecosystem structure and preserved metabolic capacity are fundamental requirements for treatment success, thereby revealing actionable targets for microbiome-modulating strategies to overcome resistance to cancer immunotherapy.

## Results

### Construction of a non-redundant metagenomic catalog for cancer immunotherapy studies

To identify general ecological principles of ICI response, we constructed a comprehensive, high-quality genomic catalog of the gut microbiome from cancer patients. We integrated 951 stool metagenomes from multiple cohorts, applying stringent quality control that retained approximately 20 billion high-quality reads (mean ± SD: 21 ± 14 million per sample) for genome assembly. This effort produced a non-redundant catalog of 3,816 metagenome-assembled genomes (MAGs) of excellent quality, with 91.9 ± 6.8% completeness and 2.13 ± 2.77% contamination (Table S1). Adhering to Genomic Standards Consortium (GSC) standards, the collection comprises 2,348 high-quality and 1,201 medium-quality MAGs, providing a foundation for taxonomic and functional profiling. Taxonomically, the catalog spans 12 phyla and is dominated by Firmicutes (2,540 MAGs, 66.6%), Actinobacteria (646 MAGs, 16.9%), and Bacteroidetes (402 MAGs, 10.5%), with representation from Proteobacteria (3.7%) and other phyla (2.3%) (Figures 1A-B). It includes 11 archaeal genomes, reflecting the phylogenetic diversity of the gut ecosystem in an oncology setting. For subsequent association analyses with clinical outcomes, we curated a homogeneous cohort of 624 samples from 11 independent studies, excluding fecal microbiota transplantation trials to avoid confounding (Table S2). Relative abundance profiling was performed using this catalog to ensure consistent genomic resolution across all samples (Table S3). This comprehensive, publicly available catalog represents a key resource for studying host-microbiome interactions in cancer immunotherapy and enables reproducible ecological analysis across cohorts.

**Figure 1.**
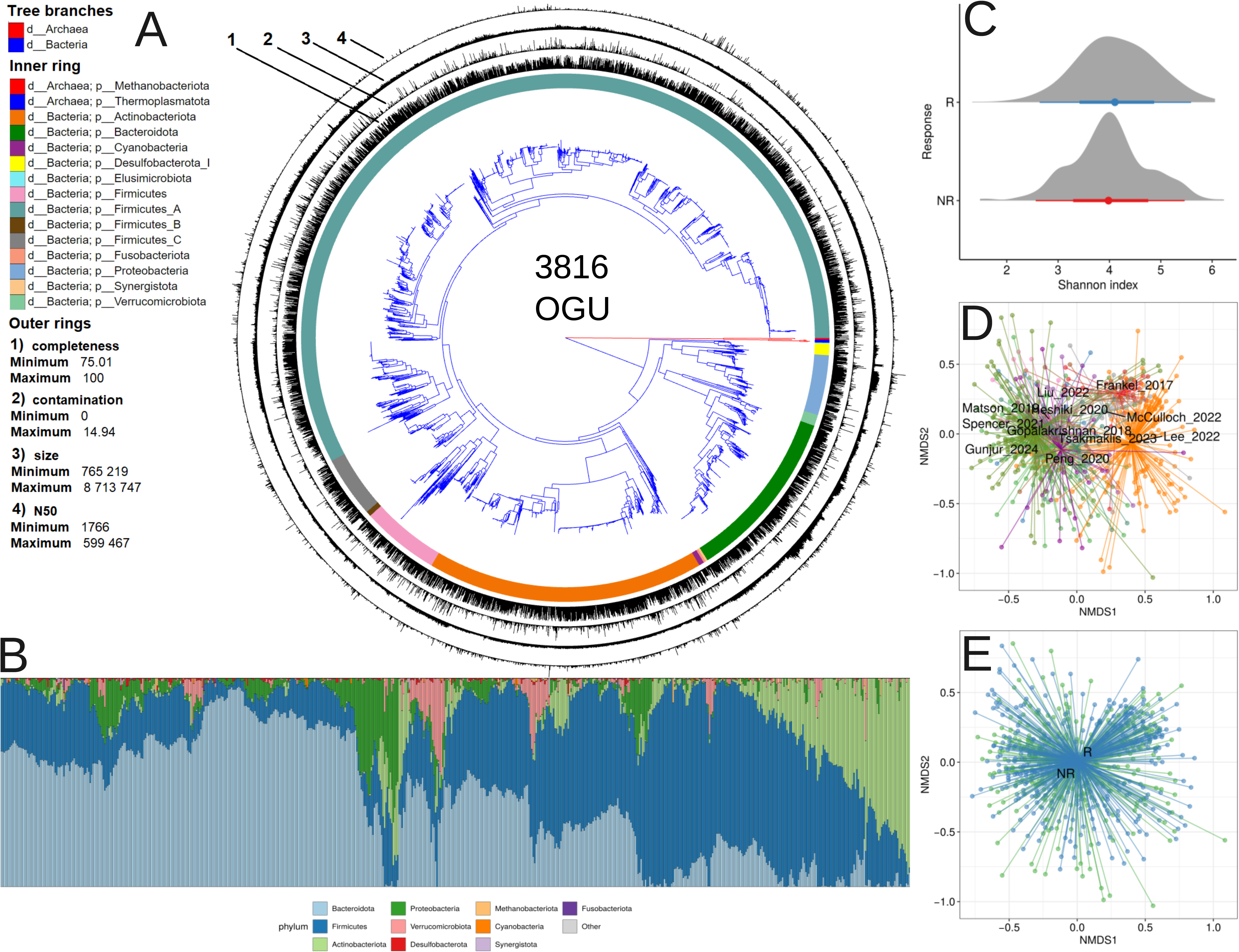
Gut microbiome catalog and ecological associations with immunotherapy outcomes. A) Phylogenetic diversity and quality assessment of the non-redundant metagenomic catalog. An approximate maximum likelihood phylogenetic tree was constructed from 3,816 MAGs using a concatenated alignment of 43 universal single-copy marker proteins (CheckM). The tree topology reveals the evolutionary relationships across the catalog. Branch colors indicate taxonomic kingdom. Concentric rings around the tree provide additional metadata: the inner colored ring displays phylum-level classification. The outer rings visualize key genome quality metrics for each operational genomic unit (OGU), allowing for direct assessment of assembly quality alongside taxonomic identity. B) Compositional profile of the analyzed cohorts. The bar plot displays the relative abundance of bacterial phyla across the 624 baseline stool samples from 11 studies, ordered by dataset. C) Comparison of microbial community richness and evenness (alpha-diversity) between responders (R) and non-responders (NR) to immunotherapy. D-E) Multivariate analysis of community structure (beta-diversity). The non-metric multidimensional scaling (NMDS) ordination based on Bray-Curtis dissimilarity reveals that sample composition clusters primarily by source dataset (D). A subtle but statistically significant separation by treatment response is detectable within this dominant technical variation (E).

### Alpha and beta diversity analysis

We first assessed whether overall microbial community richness and evenness (alpha-diversity), measured by the Shannon index, differed between responders (R) and non-responders (NR). We found no statistically significant difference between groups (linear mixed-effects model, p = 0.065; Fig. 1C). While a non-significant trend toward higher diversity was observed in responders (R: 4.13 ± 0.75; NR: 4.00 ± 0.75), the effect size was small (Cohen’s d = 0.16). As expected, the dataset source accounted for substantial variance in the model (p < 2.2e-16), justifying the use of a mixed-effects model that controlled for this technical heterogeneity. Cancer type had no significant effect (p = 0.517). These results indicate that alpha diversity, as a global ecological metric, is not a strong predictor of clinical outcome in our pan-cancer cohort, prompting us to investigate more specific community features.

To evaluate the association between gut microbiome composition and response to immune checkpoint inhibitor therapy, we performed a multivariate analysis of community structure using permutational analysis of variance (PERMANOVA). The analysis revealed that inter-study heterogeneity was a dominant source of variation, with significant dispersion differences observed across datasets (F = 4.41, p < 0.001) and cancer types (F = 14.88, p < 0.001). In contrast, no significant difference in dispersion was detected between responder groups (F = 0.04, p = 0.84), justifying the need for stratification in subsequent analyses and supporting the validity of within-stratum comparisons. After stratifying by dataset to account for study-specific effects, we analyzed the association using three distinct distance metrics. A statistically significant association between baseline microbiome structure and treatment response was first confirmed using Bray-Curtis dissimilarity (F = 1.38, R² = 0.002, p = 0.049). Furthermore, analyses using phylogenetically-informed metrics yielded stronger associations. The Weighted UniFrac analysis showed a more pronounced effect (F = 2.93, R² = 0.005, p = 0.015), and the strongest signal was detected using a distance derived from PhILR transformation (F = 4.26, R² = 0.007, p = 0.014). This ordination-based visualization aligned with the PERMANOVA results, showing primary clustering by dataset alongside a subtler, yet discernible, separation by therapeutic response (see Figure 1D-E).

### Differential abundance analysis with MaAsLin2

To identify microbial taxa associated with immunotherapy response, we performed differential abundance analysis using MaAsLin2 with mixed-effects models. This analysis was conducted on datasets Frankel_2017, Gopalakrishnan_2018, Spencer_2021, McCulloch_2022, Gunjur_2024, Heshiki_2020, and Liu_2022, adjusting for key covariates (age and gender) where applicable (Figure 2; Table S4). This analysis revealed a core set of consistent associations, as well as several context-dependent signatures linked to specific cancer types. The most robust, pan-cancer biomarker identified was the genus *Fusicatenibacter*, and specifically the species *Fusicatenibacter saccharivorans*, which was significantly enriched in R across all cohorts, even after adjusting for gender and age. In contrast, the NR state was characterized by a reproducible shift toward an aberrant profile. A key pan-cancer finding was the consistent enrichment of the phylum Proteobacteria, a hallmark of inflammation and ecosystem instability. Furthermore, the species *Parasutterella excrementihominis* emerged as a robust, pan-cancer microbial marker for non-response. Several other well-known beneficial taxa exhibited significant but context-dependent associations. The genus *Blautia* and its species *Blautia wexlerae* were positively associated with response specifically in the melanoma cohort. Similarly, *Faecalibacterium prausnitzii* was a significant marker in melanoma, but this association did not persist in the “other cancers” cohort after covariate adjustment. The genus *Gemmiger* presented a complex picture, with associations that reversed direction depending on cancer type and model adjustment; its species *Gemmiger qucibialis* was identified in opposing contexts (NR in adjusted melanoma; R in adjusted other cancers).

**Figure 2.**
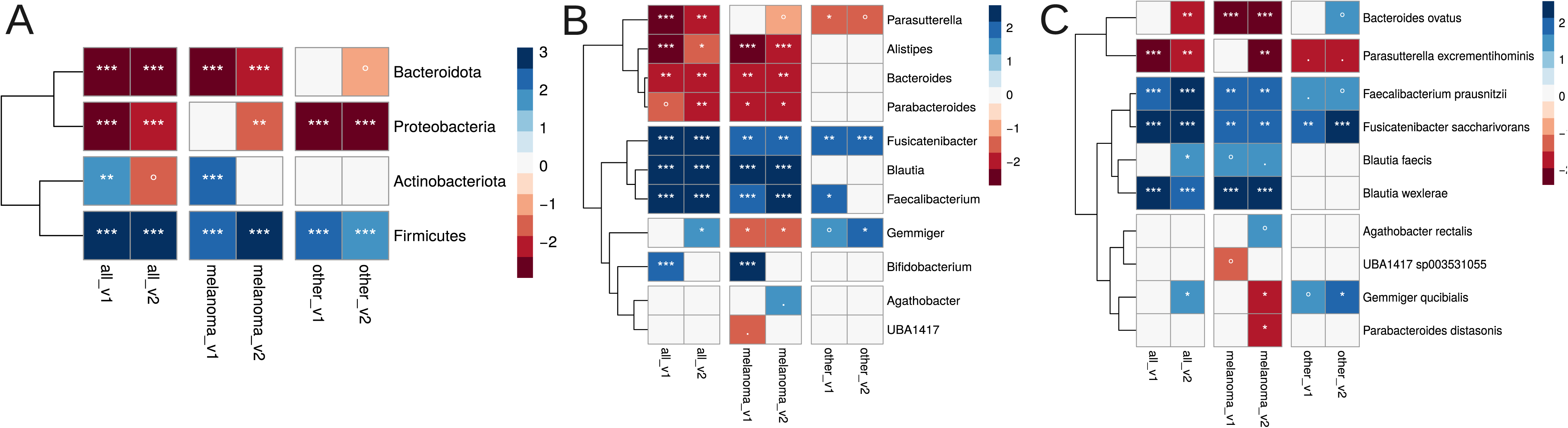
Functional enrichment analysis of taxonomic shifts identified by differential abundance analysis. Results of the functional enrichment analysis for taxonomic shifts identified as significant by the MaAsLin2 differential abundance analysis. Positive associations, where a taxon is enriched in the target condition, are shown in blue. Negative associations, where a taxon is depleted, are shown in red. Panel (A) displays significant associations at the species level, Panel (B) at the genus level, and Panel (C) at the family level. Statistical significance levels are denoted as follows: ***p < 0.001; **p < 0.01; *p < 0.05; · p < 0.1; ° p < 0.25.

### Microbial prevalence as a determinant of immunotherapy outcomes

Previous meta-analyses indicate that the most reliable microbial biomarkers of immunotherapy response are often highly prevalent, autochthonous taxa associated with intestinal health (e.g., *Faecalibacterium prausnitzii*) [Limeta et al., 2020; Zakharevich et al., 2024]. This recurring pattern suggests a broader hypothesis: a bacterium’s prevalence within a cohort may correlate with its association with a favorable outcome, as abundant resident microbes likely play key roles in maintaining immune homeostasis. In line with this, our top biomarker, the health-associated species *F. saccharivorans*, exhibited significantly higher-than-average prevalence (Wilcoxon test, p = 3.169e-12), serving as a paradigmatic example. This observation led us to hypothesize that a bacterium’s prevalence across samples in a cohort directly correlates with its association with positive immunotherapy outcomes in melanoma. We propose that highly prevalent, autochthonous microbes are evolutionarily adapted for symbiotic coexistence and play integral roles in maintaining immune homeostasis. Their abundance may therefore serve as a robust indicator of a stable gut environment conducive to treatment success. Conversely, low-prevalence bacteria represent a heterogeneous group comprising either rare autochthonous ecosystem members, which are challenging to detect consistently, or allochthonous microbes potentially originating from external sources such as food or the upper gastrointestinal tract. Standard differential abundance analysis faces substantial limitations when evaluating these low-prevalence taxa. Statistical challenges including zero-inflation (where bacteria are absent in most samples) and low abundance lead to unreliable regression coefficients with wide confidence intervals and reduced statistical power. Consequently, rare taxa are typically filtered out prior to analysis, which reduces the total number of hypotheses tested and mechanically affects the stringency of multiple comparison corrections, ultimately limiting discovery of associations with potentially informative allochthonous community members.

Initial correlation analysis of MaAsLin2 coefficients - recomputed without prevalence filtering to include all microbial taxa in the dataset - with microbial prevalence revealed a weak but statistically significant positive relationship (Spearman ρ = 0.14 ± 0.05, p < 0.001) across all analytical variants described in the previous analysis of taxonomic associations. To address the limitations of conventional approaches, we employed an alternative method based on Songbird [Morton et al., 2019], which we have successfully applied in previous studies [Olekhnovich et al., 2023; Zakharevich et al., 2024]. This approach generates regression coefficients for each bacterium without pre-filtering within each dataset independently, interpreted as log₂ fold changes while accounting for underrepresented diversity. The Songbird-based correlation analysis demonstrated stronger and more consistent effects: across all datasets, the correlation was positive in 10 of 11 datasets (Spearman ρ = 0.25 ± 0.09, adj. p < 0.001). The single exception was the Heshiki_2020 dataset (n = 11), which showed a significant negative weak correlation (Spearman ρ = −0.07, adj. p < 0.05). The robustness of this relationship was further validated using linear regression modeling, which confirmed a significant positive association between prevalence in collected dataset and Songbird regression coefficients (Estimate = 1.310e-03, p < 2e-16) without substantial confounding by dataset identity, as the dataset factor terms showed no statistically significant effects (all p > 0.05 after multiple testing consideration). Given the superior performance of Songbird in capturing the prevalence-association relationship - evidenced by both stronger correlation coefficients - we selected this approach for all subsequent analyses. This decision is further supported by Songbird’s inherent methodological advantages for compositional data, including its ability to model rare taxa thereby providing a more comprehensive and biologically relevant assessment of microbial associations with immunotherapy outcomes. Detailed correlation coefficients and p-values for each individual dataset are comprehensively documented in Table S5.

### Discovery and validation of stool microbiome-derived OGU markers predictive of immunotherapy response using Songbird

Songbird-based differential abundance analysis identified 350 marker OGUs that were significantly differentiated between R and NR groups. Among these, 149 features showed enrichment in responders, while 201 were enriched in non-responders (Table S6). We found a statistically significant correlation of moderate strength (Spearman ρ = 0.63, p-value < 2.2e-16) between the mean Songbird regression coefficient for each marker and its prevalence in the dataset. The variability of Songbird regression coefficients for these marker OGUs across different datasets reveals consistent directional effects despite inter-study heterogeneity. Analysis of log ratio values derived from metagenomic samples (Table S7) demonstrated clear separation between experimental groups, with distribution patterns visualized in Figure S1A. Mixed-effects modeling revealed a significant positive association between responder status and log ratio values (β = 1.759, p < 0.001), confirming that responders consistently exhibited higher log ratios. This indicates that a shift in the gut ecosystem toward a consortium of responder-enriched taxa is a key determinant of treatment success. The dataset variable accounted for substantial random effects variance (p < 0.001), indicating important study-specific variations while maintaining the overall significant relationship. Visual inspection of the model’s residuals confirmed that their distribution was approximately normal, supporting the validity of the statistical inference. The robustness of these findings was additionally supported by bootstrap validation, which yielded a 95% confidence interval of [0.902, 2.468] for the fixed effect estimate. Effect size analysis further substantiated the clinical relevance of this association, revealing a large practical effect (Cohen’s d = 0.82) that underscores the biological significance of the identified microbial signatures in distinguishing treatment response groups.

To rigorously evaluate the generalizability and predictive power of this log-ratio biomarker, we performed a leave-one-group-out cross-validation. Logistic regression models were trained on all but one dataset and tested on the held-out cohort. The results demonstrate stable predictive performance across diverse studies, with a mean AUC of 0.67 ± 0.13 (Figure S1B). Performance was notably strong in several independent cohorts, including Heshiki_2020 (AUC=0.90) and Gunjur_2024 (AUC=0.88). Detailed information about prediction metric scores presented in Figure S2. While the log-ratio shows stable predictive power across the cohorts we analyzed, its performance in a new, unrelated dataset could vary. This is because this parameter is based on the specific mix of microbial strains (OGUs) found in our study, which may differ in other populations which may be different from the OGU set.

### Taxonomic and ecological profiles of the identified OGU markers

We further characterized the 350 previously identified marker OGUs through taxonomic enrichment analyses (Figure 3). A key overarching finding was the establishment of Proteobacteria enrichment as a universal marker of non-response, consistently identified by both MaAsLin2 and Songbird methods. Songbird analysis recapitulated this strong signature in the NR group (NES = −1.93, adj. p = 0.007), with core species including *Enterobacter kobei*, *Escherichia coli*, and several *Citrobacter* and *Klebsiella* species. In contrast, defining a single, universal taxonomic marker of response at the phylum level proved more complex. The most notable divergence between methods concerned the phylum Bacteroidota: it was associated with the NR group in the initial MaAsLin2 analysis but emerged as significantly enriched in the R group in the present Songbird analysis (NES = 2.29, adj. p < 0.001). This discrepancy highlights the challenge of identifying a pan-cancer, phylum-level marker for success, as associations can be method-dependent and context-specific. However, the refined Songbird-based signature revealed important and consistent patterns. At finer taxonomic resolutions, the R group was robustly enriched for the genera *Bacteroides*, *Gemmiger*, and *Faecalibacterium* (NES > 2, adj. p < 0.001). Notably, these responder-linked markers predominantly belong to the most prevalent species in our dataset, reinforcing our earlier hypothesis that high-prevalence, autochthonous taxa are key players in a favorable outcome. Conversely, the NR group was characterized by the enrichment of *Prevotella* (NES = −1.78, adj. p = 0.02) and *Veillonella* (NES = −2.30, adj. p < 0.001). Collectively, these results consolidate Proteobacteria as a core, reproducible dysbiotic signature while refining the beneficial consortium to include prevalent Bacteroidota and other high-abundance genera, as more accurately captured by the Songbird framework.

**Figure 3.**
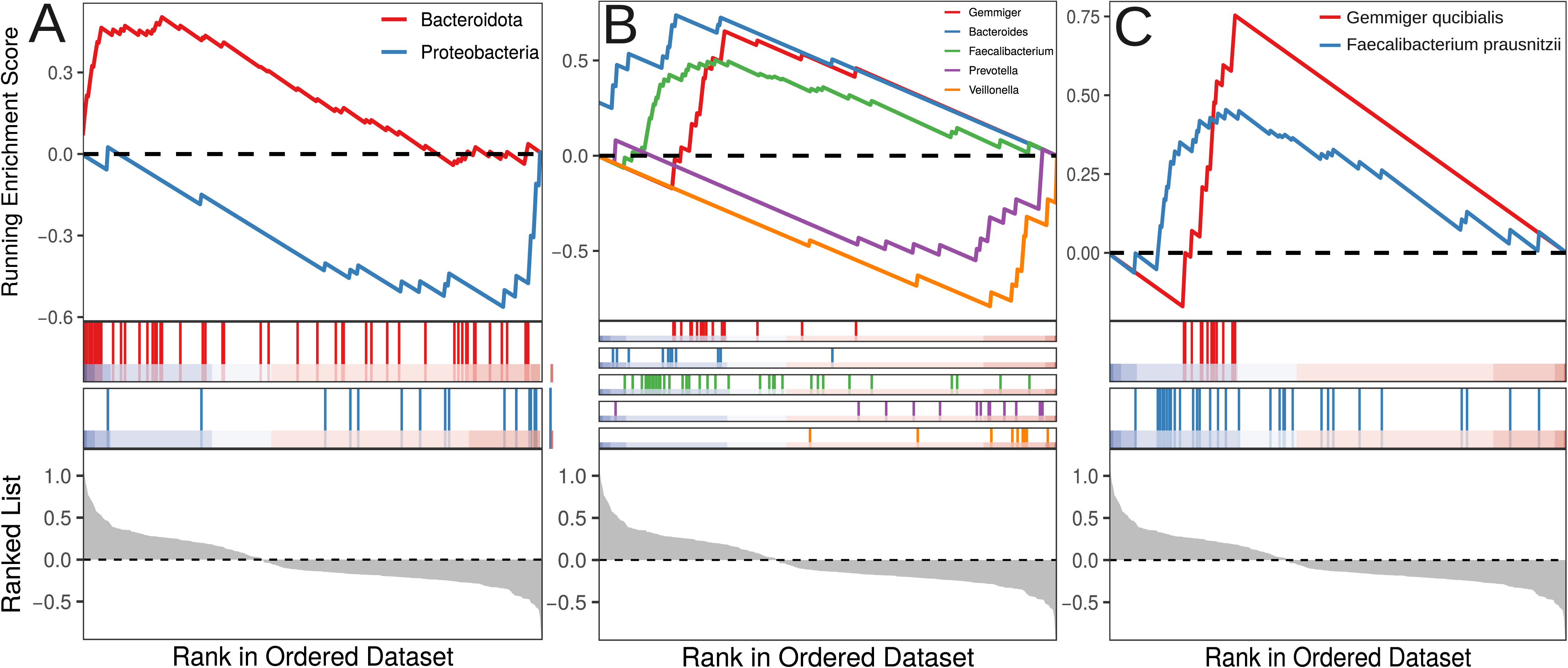
Taxonomic enrichment analysis characterizing the 350 marker OGUs identified by Songbird. The results of a Gene Set Enrichment Analysis (GSEA) are presented to evaluate the coordinated enrichment patterns of the identified marker Operational Genomic Units (OGUs) across phylogenetically ranked lists. The top panel in each subfigure displays the running Enrichment Score (ES) curve, where individual lines are colored according to their corresponding taxonomy. The position of the maximum ES (peak) indicates where the marker set is most concentrated within the ranked list; a positive score denotes enrichment at the top of the ranking (associated with the R group), while a negative score denotes enrichment at the bottom (associated with the NR group). The middle panel shows the distribution of the marker OGUs (vertical bars) across the same ranked list of all taxa, indicating where members of the significant set appear. The bottom panel illustrates the value of the ranking metric for each taxon in the list, which forms the basis for the ranking on the x-axis. This integrated visualization identifies biologically coherent taxonomic groups whose members are non-randomly distributed towards the extremes of the phenotypic ranking. The analysis is shown at distinct taxonomic resolutions: (A) phylum level, (B) genus level, and (C) species level.

To investigate exogenous origins, we compared our 3,816 OGUs to MAGs from non-intestinal body sites and food (Figure 4). We identified 293 OGUs clustering with non-intestinal MAGs and 91 with food-derived MAGs at the species level (95% ANI; Table S8). Strikingly, OGUs associated with non-responders (NR) were significantly enriched within these food-derived (NES = −2, *p* = 0.001) and oral-derived (NES = −2.60, *p* = 5.18e-7) clusters. This pattern indicates an aberrant microbial state in non-responders, characterized by an increased prevalence of core species we define as food-borne opportunists (e.g., *Citrobacter freundii* and *Klebsiella michiganensis*, both Proteobacteria) and oral commensals (e.g., *Veillonella parvula* and *Anaeroglobus micronuciformis*).

**Figure 4.**
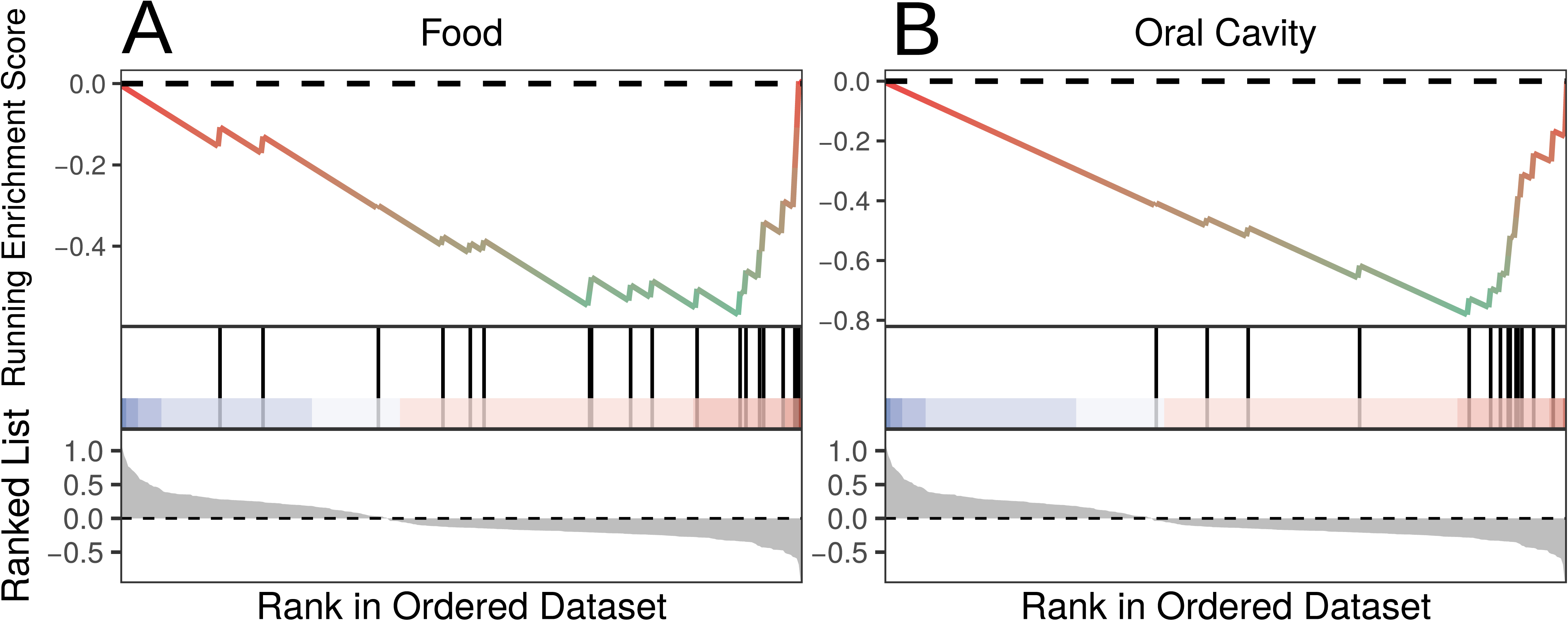
Enrichment analysis of marker OGUs across non-intestinal body sites and food origins. The enrichment analysis is presented to evaluate the distribution and enrichment of the identified marker OGUs across ranked lists of microbial taxa associated with specific bodily or dietary sources. The top panel in each subfigure displays the running enrichment score (ES) curve, where individual lines are colored according to their assigned source category. A positive ES peak indicates the marker OGUs are significantly concentrated among taxa associated with that source and enriched in responders to immunotherapy, whereas a negative score indicates concentration among taxa associated with that source and enriched in non-responders to immunotherapy. The middle panel shows the position of the marker OGUs (vertical bars) within the ranked list of all source-associated taxa. The bottom panel illustrates the ranking metric value for each taxon, defining the order on the x-axis. This visualization identifies whether the marker OGUs are collectively enriched for microbial signatures typical of specific non-intestinal body habitats or dietary origins. The analysis is stratified by source type: (A) food origin, and (B) oral cavity origin.

### Functional profiles of the identified OGU markers

Analysis of the carbohydrate-active enzyme (CAZy) repertoire revealed a distinct functional capacity associated with positive immunotherapy outcome. Microbial markers linked to responders contained a significantly higher frequency of CAZy genes than non-responder-associated markers (Wilcoxon rank sum test, W = 19,802, p = 1.24e-07; Figure 5A). Furthermore, Fisher’s exact test identified 27 CAZy families significantly enriched in the responder group compared to only 3 in non-responders (logFC > |1|, adj. p < 0.05; Table S9, Figure 5B). The enriched CAZy profile in responders indicates specialized adaptation for breakdown of complex dietary and host-derived carbohydrates. This includes polysaccharide lyases PL1, PL9, PL10, PL15 and PL33 along with glycoside hydrolases GH26, GH28, GH115 and GH130 involved in plant cell wall decomposition. Notably, we observed enrichment of mucin-degrading enzymes GH89, GH125, GH129 and GH171, which target key components of intestinal mucin glycans. The profile was complemented by carbohydrate esterases CE1, CE2, CE4, CE7, CE12 and CE17, alongside carbohydrate-binding modules CBM13, CBM32 and CBM67, collectively indicating sophisticated capability for complex carbohydrate breakdown. Enrichment analysis further confirmed the overrepresentation of glycoside hydrolase families in responders (NES = 1.35, p = 0.02; Figure 5C). The core enrichment comprised 47 GH families encompassing hemicellulases GH30, GH43, GH51, GH115, GH120, GH26, GH27 and GH113; involved to pectin degradation GH28, GH78, GH88 and GH105; and intestinal adaptation specialists GH29, GH95, GH33, GH89, GH125, GH129, GH2, GH20 and GH35. This functional arsenal also included enzymes for bacterial biofilm degradation GH32, GH66 and GH144, peptidoglycan cleavage GH23, GH24 and GH25, and chitin degradation GH18 and GH20, indicating superior competitive capacity within the gut ecosystem (Table S10). Collectively, these results demonstrate that responder-associated microbes represent a highly efficient and competitive community for carbohydrate processing, strongly adapted to the rich and competitive environment of the mammalian colon. Their ability to utilize both dietary plant fibers and host mucin provides metabolic flexibility, while their arsenal of enzymes targeting bacterial cell walls and biofilms suggests enhanced competitive fitness.

**Figure 5.**
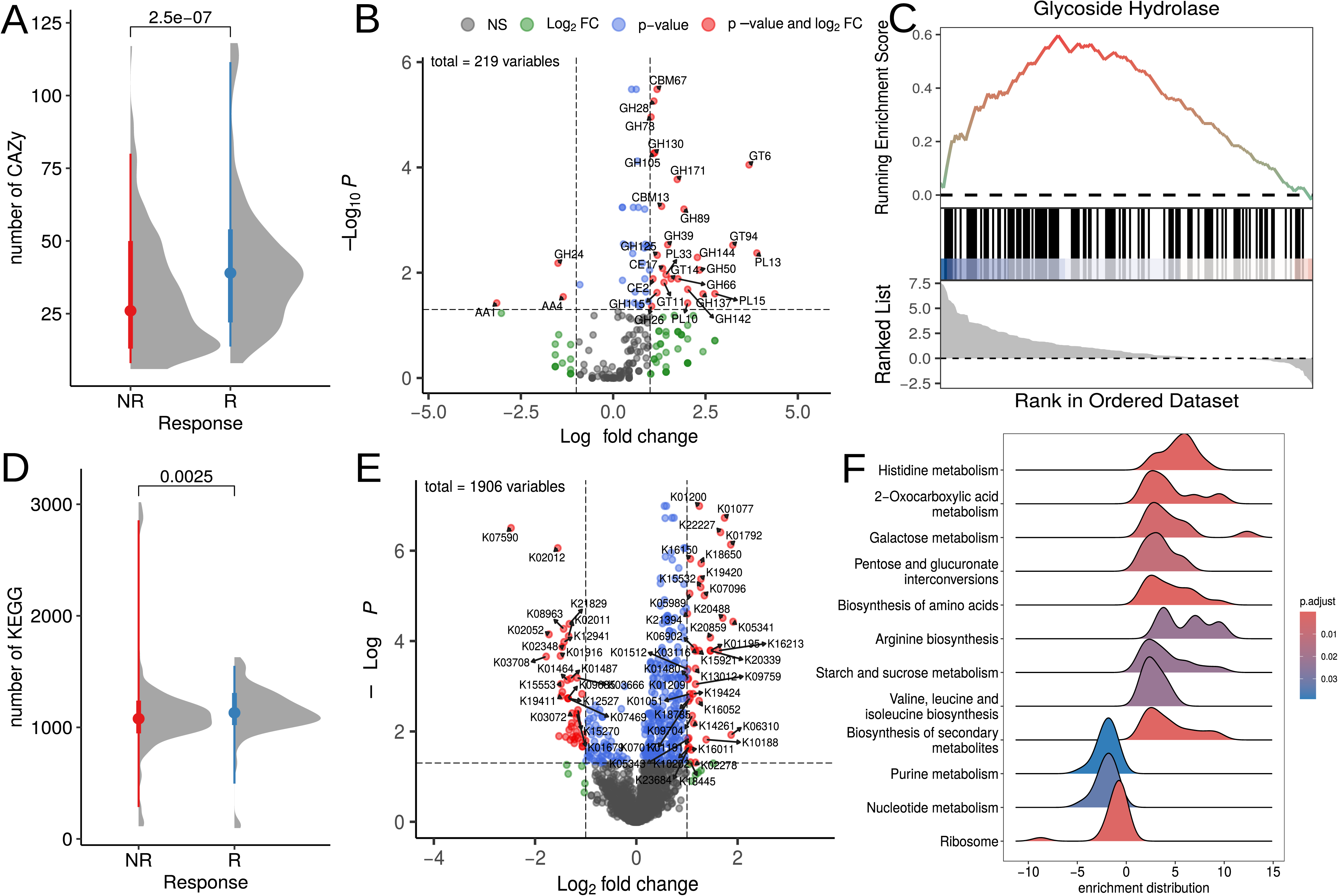
Functional profiling of immunotherapy response-associated marker OGUs. Analysis of the functional potential encoded by the marker OGUs associated with responders and non-responders to immunotherapy. (A-C) Carbohydrate-active enzyme (CAZy) repertoire analysis. (A) The overall frequency of CAZy genes identified within the marker OGUs for each response group. (B) Specific CAZy families significantly enriched in the marker OGUs of responders (positive values) and non-responders (negative values). (C) Enrichment analysis results highlighting glycoside hydrolase families that are coordinately enriched across the ranked list of all OGUs, with positive scores indicating enrichment in responders and negative scores in non-responders. (D-F) Functional profile based on KEGG orthology. (D) The total number of unique KEGG orthology (KO) groups present in the marker OGUs of each response group. (E) Specific KO groups showing significant differential enrichment between responders and non-responders. (F) Pathway-level GSEA results depicting KEGG pathways significantly upregulated (positive enrichment scores) in the responder-associated marker OGUs.

Functional annotation based on KEGG orthology revealed a distinct functional potential between the R and NR groups. A significantly higher overall number of KEGG orthology groups was observed in the R group compared to the NR group (Wilcoxon rank sum test, W = 17810, p-value = 0.001227; Figure 5D). Differential abundance analysis using a Fisher’s exact test identified 43 significantly enriched genes in the R group and 42 in the NR group (logFC > |1|, adj. p < 0.05; Table S11, Figure 5E). Subsequent gene-set enrichment analysis (GSEA; NES > |1|, adj. p < 0.05; Table S12, Figure 5F) further delineated this functional divergence. The R group exhibited a pronounced upregulation of pathways central to metabolic processing, specifically encompassing ko00340 (Histidine metabolism), ko01210 (2-Oxocarboxylic acid metabolism), ko00052 (Galactose metabolism), ko00040 (Pentose and glucuronate interconversions), ko01230 (Biosynthesis of amino acids), ko00220 (Arginine biosynthesis), ko00500 (Starch and sucrose metabolism), ko00290 (Valine, leucine and isoleucine biosynthesis), and ko01110 (Biosynthesis of secondary metabolites). In contrast, the NR group was characterized by a significant enrichment of pathways fundamental to genetic information processing and cellular maintenance, namely ko00230 (Purine metabolism), ko01232 (Nucleotide metabolism), and ko03010 (Ribosome). This functional signature suggests that the R-group microbiota is primed for active degradation of complex dietary substrates and biosynthesis of essential metabolites, whereas the NR community exhibits a profile more focused on core cellular maintenance and replication.

## Discussion

The past decade has witnessed an exponential growth in human microbiome data, leading to the identification of numerous taxonomic associations with various pathologies. Meta-analyses consistently demonstrate reproducible, almost archetypal patterns of dysbiosis: a depletion of symbiotic microbes such as *Faecalibacterium*, *Roseburia*, and *Blautia* - highly prevalent components of a healthy microbiome - across a wide spectrum of diseases, from inflammatory bowel pathologies to metabolic syndromes and differential responses to cancer immunotherapy [Pascal et al., 2017; Sun et al., 2024; Romano et al., 2025; Lin et al., 2025]. Concurrently, representatives of Proteobacteria (including *Escherichia coli*) [Pascal et al., 2017; Akiyama et al., 2024], as well as taxa characteristic of the oral microbiota, such as *Streptococcus salivarius*, *Bifidobacterium dentium*, and *Veillonella parvula* [Dubinkina et al., 2017; Brennan et al., 2019; Rojas-Tapias et al., 2022; Stagaman et al., 2024; Li et al., 2024; Manghi et al., 2025], have emerged as negative biomarkers in various contexts. Progress in the field has not been limited to establishing correlations. The causal role of specific microbial agents, for instance, certain strains of *Escherichia coli* in the pathogenesis of inflammatory bowel disease and colorectal cancer, has been experimentally proven in model systems [Pleguezuelos-Manzano et al., 2020; Viladomiu et al., 2021]. Nevertheless, the integration of these data often fosters a simplified dualistic view: “*Faecalibacterium* is good, Proteobacteria is bad.” We posit that these classic, reproducible taxonomic markers are more a consequence than a cause, reflecting more fundamental ecological shifts within the microbial community. These shifts find their conceptual reflection in the Anna Karenina principle for microbiomes [Zaneveld et al., 2017]. According to this principle, healthy microbial communities are characterized by a relatively uniform and stable structure (“all happy families are alike”), whereas in dysbiosis, their composition becomes highly variable and idiosyncratic (“each unhappy family is unhappy in its own way”).

The Anna Karenina principle for microbiomes finds direct confirmation in the context of immuno-oncology within this study. We propose an ecological interpretation of this phenomenon, based on the identified positive correlation between a taxon’s prevalence in the population and its association with a positive response to therapy. The obtained data align with the hypothesis that the “happy uniformity” of responders is driven by the dominance of a highly prevalent, evolutionarily stable core of autochthonous symbionts. This core is characterized by a conserved functional profile that supports the host’s metabolic needs. This conclusion receives direct molecular validation from our data: patients who responded to therapy exhibit enrichment of their microbial community in genes for carbohydrate-active enzymes (CAZy), particularly glycoside hydrolases linked to plant polysaccharide metabolism, activation of KEGG pathways for amino acid biosynthesis, and an overall heightened metabolic potential. The evolutionary fitness of this symbiotic core is further evidenced by a specific profile of mucin-degrading CAZy families (GH89, GH125, GH129, GH171). These enzymes facilitate shallow, controlled degradation of mucin, a trait characteristic of adapted commensals rather than aggressive colonizers.

It can be hypothesized that at a global level, a healthy microbiome converges toward a similar functional optimum among individuals within a population. Factors such as an imbalanced diet, stress, and antibiotic use disrupt community assembly processes, leading to reduced functional resilience and, consequently, a diminished ability to support an adequate immune response. This perspective finds direct support in clinical data: for instance, high intake of plant-based, fiber-rich foods is associated with improved immunotherapy outcomes and promotes the formation of a favorable microbiome in model experiments, whereas direct probiotic administration does not yield a comparable effect [Spencer et al., 2021]. This aligns with the idea that success is determined by creating conditions for the self-organization of the autochthonous community, rather than by introducing isolated components into it. This interpretation also helps explain the widely observed inconsistency in taxonomic signatures across differential abundance analyses and the problem of high sparsity in abundance matrices characterized by numerous zero values. We propose that this is a direct consequence of a fundamental ecological principle of community assembly. For each individual, regardless of geographic or climatic zone, it is ecologically possible to form a unique, yet functionally competent assembly of symbionts, adapted to local conditions including diet shaped by sociocultural norms and the microbial landscape of their immediate environment. Thus, a healthy, evolutionarily optimal microbiome represents a highly individualized combination of taxa, which logically results in low concordance at the species level when comparing different cohorts. This underscores that the key properties of the ecosystem - stability and the ability to support host homeostasis - are determined not by a specific taxonomic composition, but by the preservation of a functional core and the community’s capacity for self-organization within given ecological constraints. We suggest that this is underpinned by a universal principle governing the assembly of complex microbiological systems: the evolution of a microbial ecosystem tends toward a functional optimum that can be achieved by multiple different taxonomic configurations depending on the local context [Burcham et al., 2024].

Conversely, a dysbiotic state - whether due to poor diet, antibiotic use, or disease - represents the collapse of this complex system. It is succeeded by a “selfish” ecological strategy. In the newly vacated niche, competitive advantage shifts to taxa focused on maximizing the use of simple resources for their own replication (which may be reflected in the enrichment of nucleotide metabolism pathways), rather than on producing system-wide benefits. In the clinical context, this state reflects the ecosystem’s loss of colonization resistance and its inability to provide the immunomodulatory tone necessary for an effective therapeutic response. Therefore, the key predictor of outcome is not merely the presence of individual “good” genes or taxa, but the preservation of the ecosystem’s intrinsic capacity to maintain a complex, distributed metabolism shared among different species. From this perspective, restoring clinical response may be linked not to specific bacterial replacement therapy, but to creating conditions that allow the reassembly of the autochthonous functional core and the restoration of cooperative interactions within the community. It is worth noting that research into fecal microbiota transplantation from responding patients to non-responding patients substantiates this thesis [Baruch et al., 2021; Davar et al., 2021]. Responding patients likely retained a larger portion of the communal core, which compensated for the parts lost in non-responders. Furthermore, the thesis of a universally beneficial microbiome, not only in the context of immunotherapy, is supported by the effectiveness of FMT from healthy donors to improve immunotherapy efficacy [Routy et al., 2023].

However, the strength of the identified association between a taxon’s prevalence in the population and a favorable outcome is moderate. This likely reflects a fundamental heterogeneity in the composition within the “tail” of the species abundance distribution. This region harbors two ecologically distinct groups: true low-abundance autochthonous symbionts and transient allochthonous species originating from proximal parts of the gastrointestinal tract (e.g., the oral cavity) or introduced with food. Confirmation of this hypothesis was obtained through an analysis based on clustering operational genomic units (OGUs) from the assembled catalog with sets of genomes from various human biotopes and food products. This finding is supported by our analysis, which demonstrates that taxa associated with a lack of response to therapy are enriched for OGUs that cluster together with metagenome-assembled genomes (MAGs) from the oral microbiota and food. This quantitatively confirms the mixing of heterogeneous ecological signals in the low-abundance region. The interpretation of these data is complicated by the fundamentally compositional nature of metagenomic data [Gloor et al., 2017]. This observation aligns with our previous results from analyzing gut colonization after fecal microbiota transplantation [Olekhnovich et al., 2021]. In that work, we showed that the probability of successful colonization of a taxon in a recipient directly correlates with its initial abundance in the donor: highly abundant taxa predictably persist or engraft, whereas for low-abundance taxa, predicting the outcome becomes statistically problematic. We hypothesized that this is precisely because the “tail” of the donor’s abundance distribution is a mixture of true low-abundance residents and transient allochthonous microbes passing through the gut, making them ephemeral and unreliable for targeted transmission. The observed relative enrichment of oral taxa in patients who did not respond to immunotherapy can be explained by two complementary hypotheses. First, it could be an artifact reflecting a sharp decrease in the absolute biomass of the autochthonous core. In this case, the relative increase in the proportion of oral taxa is not a consequence of their true expansion but a result of compositional distortion against the background of the “disappearance” of dominant symbionts, consistent with data on the increase in their relative abundance when overall microbial load decreases [Liao et al., 2024]. Second, it may reflect a true ecological colonization process. Reduced butyrate production, associated with decreased diversity and the loss of key symbionts, leads to impaired colonocyte nutrition, thinning of the mucin layer, and consequently, weakened colonization resistance. This creates conditions for the adhesion and persistence of commensals or opportunistic pathogens from other niches (oral, dietary), which potentiates the formation of a pro-inflammatory microenvironment and systemic immune dysregulation.

## Limitations

While this study, being primarily theoretical in nature, provides a comprehensive ecological framework for understanding microbiome-mediated response to immunotherapy within the field of host-associated microbial ecology, several limitations should be considered when interpreting the results.

Our analysis identifies robust associations between microbial community features, their functional potential, and clinical outcomes. However, the observational design of the included cohorts precludes definitive causal conclusions. Although we integrate evidence from interventional studies and mechanistic experiments from the literature to support a plausible causal pathway, future randomized controlled trials are needed to confirm that modulating the identified ecological state directly enhances treatment efficacy.

Like all metagenomic studies, our analysis is based on relative abundance data, which is inherently compositional. This limits direct inference about absolute changes in microbial biomass. The observed shifts, such as the relative increase in oral taxa in non-responders, could stem from a true expansion of these groups, a severe decrease in the autochthonous core biomass, or a combination of both. While we reference studies linking low microbial load to dysbiosis [Liao et al., 2024], concurrent measurements of absolute bacterial load would be required to disentangle these effects fully.

The functional profiling is based on genomic potential inferred from MAGs and gene catalogs. This reflects the community’s functional capacity but not its actual activity or metabolite production in vivo. Transcriptomic, proteomic, or metabolomic analyses of patient samples are necessary to validate that the predicted metabolic pathways, such as butyrate production or mucin degradation, are actively contributing to the host-microbe dialogue in responders.

The central concepts of “ecosystem stability,” “functional core,” and “self-organization capacity” are interpretative frameworks derived from multivariate patterns. We propose specific, measurable proxies for them (e.g., prevalence, functional gene abundance, log-ratio values). However, these are partial proxies. Developing integrative, quantitative indices for ecosystem resilience and functional coherence remains a critical methodological frontier for the field.

Reference-based clustering of exogenous taxa. The identification of oral- and food-derived taxa in our catalog relies on clustering our OGU with MAGs from external reference databases at the species level (95% ANI). While this provides strong evidence for exogenous origins, it remains an approximation. Strain-level differences and ecological adaptations that occur after a microbe enters a new niche (e.g., the gut) may not be captured. Future studies employing concurrent multi-site sampling (e.g., oral and gut microbiota from the same individuals) and higher-resolution strain-tracking methods are needed to precisely map the routes of colonization, quantify the actual establishment of exogenous taxa, and understand their functional evolution in the new environment.

## Conclusions

The study created a comprehensive catalog of gut microbiome genomes, deepening our understanding of its role in cancer immunotherapy. It was established that a successful treatment response is associated not with overall species diversity, but with the specific structure and functional state of the microbial community. Specific taxonomic markers were identified, among which bacteria characteristic of dysbiosis play a key negative role, while autochthonous symbiotic species play a positive role. Methodologically, it was confirmed that a high prevalence of a taxon in the population correlates with its beneficial effect on treatment outcome. The functional profile of the microbiota of responders reflects its high metabolic activity and adaptation to the intestinal environment, in contrast to the community of non-responders. Thus, this study justifies a shift from the search for single markers to a comprehensive assessment of the ecological and functional status of the microbiota for the development of new diagnostic and therapeutic approaches in oncology.

## Materials and methods

### Collection data

To create a comprehensive data resource for characterizing the gut microbial diversity in melanoma patients undergoing immunotherapy, we reconstructed a catalog of metagenome-assembled genomes (MAGs) from 951 stool metagenomes across 14 public studies. This collection includes ten melanoma-specific datasets alongside four datasets from other cancer types. Additionally, it incorporates metagenomic data from studies investigating fecal microbiota transplantation (FMT) in melanoma immunotherapy. The complete dataset summary is presented in Table 1.

**Table 1.** Summary table of characteristics of used data.

| Dataset | Immunotherapy type | Cancer type | FMT | Samples number | Sequencing platform (sequence length, bp) | Total number of reads |
| --- | --- | --- | --- | --- | --- | --- |
| Frankel et al. 2017 [2] | anti-PD-1/<br>anti-CTLA4 | Melanoma | False | 39 | Illumina HiSeq 2000 (100) | 1,688,904,099 |
| Gopalakrishnan et al. 2018 [4] | anti-PD-1 | Melanoma | False | 22 | Illumina HiSeq 2000 (100) | 398,245,066 |
| Matson et al. 2018 [5] | anti-PD-1 | Melanoma | False | 38 | Illumina NextSeq 500 (150) | 1,522,421,744 |
| Baruch et al. 2021 [6] | anti-PD-1 | Melanoma | True | 42 | Illumina HiSeq X Ten (150) | 737,390,881 |
| Davar et al. 2021 [7] | anti-PD-1 | Melanoma | True | 214 | Illumina NovaSeq 6000 (150) | 2,422,776,370 |
| Spencer et al. 2021 [8] | anti-PD-1 | Melanoma | False | 134 | Illumina HiSeq 2000/X (100-150) | 2,028,356,126 |
| Lee et al. 2022 [9] | anti-PD-1/<br>anti-CTLA4 | Melanoma | False | 164 | Illumina NovaSeq 6000 (150) | 4,219,730,439 |
| McCulloch et al. 2022 [10] | anti-PD-1 | Melanoma | False | 27 | Illumina NovaSeq 6000 (150) | 922,889,774 |
| Tsakmaklis et al. 2023 [11] | anti-PD-1/<br>anti-CTLA4 | Melanoma | False | 29 | Illumina NovaSeq 6000 (150) | 794,124,064 |
| Routy et al. 2023 [12] | anti-PD-1 | Melanoma | True | 71 | Illumina NovaSeq 6000 (150) | 1,565,577,436 |
| Peng et al. 2020 [3] | anti-PD-1 | Cancer of part of the gastrointestinal tract (GIT) | False | 40 | Illumina NovaSeq 6000 (150) | 1,015,338,636 |
| Liu et al. 2022 [13] | anti-PD-1 | Non-small cell lung cancer (NSCLC) | False | 14 | Illumina HiSeq 4000 (150) | 409,403,683 |
| Heshiki et al. 2020 [14] | anti-PD-1 | NSCLC, cancer of part of the GIT, breast cancer, ovarian cancer | False | 11 | Illumina HiSeq 1500 (100) | 391,957,976 |
| Gunjur et al. 2024 [15] | anti-PD-1/<br>anti-CTLA4 | Cancer of part of the GIT, other rare cancers | False | 106 | Illumina NovaSeq 6000 (150) | 2,227,048,564 |

The further metagenomic analysis incorporated 624 stool metagenomes obtained from 11 collected datasets (Table 1, FMT True) excluding fecal transplant data (Table 1, FMT False). Patients were stratified by immunotherapy response into two groups: responders (R group, n = 362; 58.2%) and non-responders (NR group, n = 262; 41.8%). Response assessment followed RECIST 1.1, with the R group including patients showing complete response (CR), partial response (PR), or stable disease (SD) at 6-month follow-up, while the NR group comprised exclusively progressive disease (PD) cases. The study cohort received various immunotherapy regimens, including anti-PD1, anti-CTLA4, or combination therapies. Cancer type distribution revealed melanoma predominance (n = 456; 72.7%) [Frankel et al. 2017; Gopalakrishnan et al. 2018; Matson et al. 2018; Spencer et al. 2021; Lee et al. 2022; McCulloch et al. 2022; Tsakmaklis et al. 2023], followed by gastrointestinal (GI) cancers (n = 82; 13.1%) [Peng et al. 2020; Heshiki et al. 2020; Gunjur et al. 2024], non-small cell lung cancer (n = 15; 2.4%) [Liu et al. 2022; Heshiki et al. 2020], breast cancer (n = 4; 0.6%) [Heshiki et al. 2020], ovarian cancer (n = 2; 0.3%) [Heshiki et al. 2020], and other malignancies (n = 68; 10.8%) [Gunjur et al. 2024]. All samples were collected prior to treatment initiation to evaluate baseline microbiota status. The datasets are plotted on a world map as shown in Figure 6.

**Figure 6.**
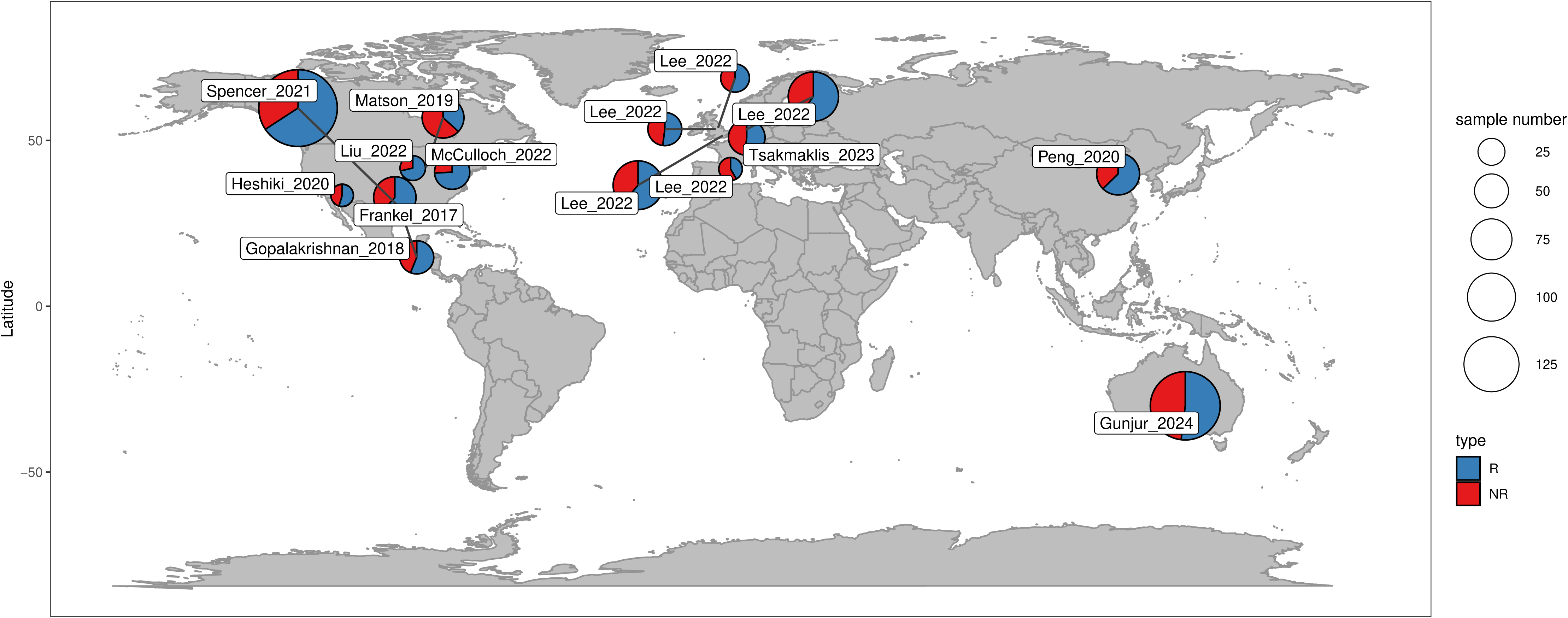
Global distribution of collected samples (n = 624) across countries used for metagenomic analysis. Circle size represents sample count per region, while color indicates the proportion of patients R - responsive (or NR - non-responsive) to immunotherapy.

### Data acquisition and preprocessing

All 951 stool metagenomic samples from 14 published studies were downloaded from the NCBI/EBI databases using Kingfisher v0.4.1 [Woodcroft, 2022]. All downloaded raw reads were preprocessed with fastp (v0.23.4) [Chen et al., 2018] with following parameters “-detect_adapter_for_pe -overrepresentation_analysis -correction -dup_calc_accuracy 6 -average_qual 30”, human DNA sequences were filtered with HISAT2 (v2.2.1) [Kim et al., 2019]. Quality control was performed using FastQC (v0.12.1) [Andrews et al., 2023] and MultiQC (v1.18) [Ewels et al., 2016].

### MAGs catalog assembly, taxonomic profiling and analysis

Metagenomic assembly was performed using MEGAHIT v1.2.9 [Li et al., 2015], retaining contigs longer than 1000 bp. Filtered contigs were aligned to metagenomic reads using HISAT2 v2.2 [Kim et al., 2019]. Binning was conducted in two stages: (1) initial binning with MetaBAT2 v2.12.1 [Kang et al., 2019], MaxBin2 v2.2.7 [Wu et al., 2016], and Semibin2 v2.1.0 [Pan et al., 2023], followed by (2) refinement with DAS Tool v1.1.7 to improve bin quality [Sieber et al., 2018]. Bins were quality-checked with CheckM v1.2.3 [Parks et al., 2015] and dereplicated at 98% nucleotide identity using dRep v3.4.5 [Olm et al., 2017] to generate operational genomic units (OGUs) by analogy with operational taxonomic units (OTUs). Zhu et al. (2022) first introduced this term, but our approach to representing OGUs offers a key advantage: OGUs are generated directly from the analyzed data, rather than through a mapping process. This more faithfully corresponds to the core concept of the “operational unit”. For the first time this approach was applied in the work of Zakharevich et. al. (2024). Taxonomic annotation of bins was performed using GTDB-Tk v2.1.1 [Chaumeil et al., 2022] with the GTDB r207 database [Parks et al., 2022]. Additional data processing utilized Samtools v1.17 [Li et al., 2009], BEDTools v2.31.0 [Quinlan et al., 2010], and BBMap v39.06 [Bushnell et al., 2014]. Finally, OGU abundance profiles were generated by mapping reads to the OGU catalog using HISAT2 v2.2, followed by processing with InStrain v1.9.0 [Olm et al., 2021].

### Alpha and beta diversity estimation

Differences in Shannon index between responders and non-responders were tested using a linear mixed-effects model implemented in R with the lmerTest package v3.1.3 [Kuznetsova et al., 2017]. Treatment response and cancer type were included as fixed effects (alpha diversity ∼ response + cancer type + (1 | dataset)). To account for systematic variation between source studies, the dataset was included as a random effect. The variance explained by the random effect (dataset) was highly significant (p < 2.2e-16), validating the model design. Assumptions of normality and homoscedasticity of residuals were verified graphically. Effect size for the group comparison was calculated as Cohen’s d using the cohens_d function from rstatix package v0.7.3.

To assess the association between gut microbiome composition and response to immune checkpoint inhibitor therapy, we performed a multivariate analysis of beta-diversity. Prior to formal hypothesis testing, the homogeneity of multivariate dispersions was evaluated using a permutational analysis of multivariate dispersions test with the betadisper function, based on 9999 permutations. This test assessed the key assumption of comparable within-group variation for the factors of dataset, cancer type, and response group. The main association between community structure and clinical outcome was then tested using a stratified permutational multivariate analysis of variance (PERMANOVA) with 999 permutations. To control for dominant inter-study heterogeneity, the PERMANOVA model was applied within strata defined by the source dataset, and the overall significance was determined by combining stratum-specific results (ogu_table ∼ response, strata = dataset). The analysis employed three complementary distance metrics to capture different ecological dimensions. Bray-Curtis dissimilarity was used as an abundance-based metric. Weighted UniFrac distance was calculated using the phyloseq package v1.52.0 [McMurdie et al., 2014]. Additionally, a phylogenetically-informed Euclidean distance was generated by first applying the PhILR (Phylogenetic Isometric Log-Ratio) [Silverman et al., 2017] transform to address data compositionality and create phylogenetically-aware balances using philr package v1.34.0. For visualization, community patterns were displayed using non-metric multidimensional scaling (NMDS) ordination based on Bray-Curtis dissimilarity. All analyses were conducted in R version v4.4.2. Multivariate procedures including betadisper, PERMANOVA and NMDS were performed using the vegan package v2.7.2 [https://github.com/vegandevs/vegan].

### MaAsLin2-differented taxonomic analysis framework

To identify taxa differentially associated with treatment response, we implemented a tiered analytical strategy designed to ensure robustness against dataset imbalances and confounding variables. Our cohort was characterized by a substantial imbalance between melanoma (n = 453) and other cancer types (n = 171), necessitating separate validation of markers within each subgroup. Our primary analysis employed MaAsLin2 v1.22.0 [Mallick et al., 2021] within a mixed-effects model framework, with immunotherapy response or cancer type in combined analysis as a fixed effect and dataset included as a random effect. We conducted parallel analyses across three data partitions: the full cohort (n = 624), melanoma-only subset (n = 453), and non-melanoma cancers (n = 171). To control for potential demographic confounders, we performed a secondary analysis on a restricted sample with available metadata (n = 353, included 7 datasets - Frankel_2017, Gopalakrishnan_2018, Spencer_2021, McCulloch_2022, Gunjur_2024, Heshiki_2020, Liu_2022), incorporating gender and age as additional fixed effects while maintaining the same random effects structure. To overcome the limited statistical power of individual taxon analyses in sparse microbiome data, we employed a systems biology approach. We aggregated MaAsLin2 results using gene set enrichment analysis (GSEA) across taxonomic hierarchies with the clusterProfiler package v4.16.0 [Xu et al., 2024], ranking features by their MaAsLin2 regression coefficients with threshold FDR < 0.1. This approach amplifies coherent biological signals by accumulating weak but consistent effects across related taxa, thereby enhancing detection of biologically meaningful patterns that might be lost in conventional single-taxon analyses. To comprehensively assess the relationship between microbial prevalence and treatment association, we conducted a correlation analysis between OGU prevalence and MaAsLin2-derived coefficients. For this specific analysis, MaAsLin2 was run without prevalence or abundance filtering (min_abundance = 0, min_prevalence = 0) to include the complete taxonomic spectrum. The association was quantified using Spearman’s rank correlation, with statistical significance determined using the cor.test function in R, followed by Benjamini-Hochberg multiple testing correction via the p.adj function.

### Determination of OGUs associated with immunotherapy outcomes using Songbird

In this study, we intentionally deviated from conventional hypothesis-driven frameworks targeting individual microbial taxa to circumvent limitations intrinsic to metagenomic data analysis. Instead, we implemented a differential ranking approach to prioritize features based on their association strength with clinical response to immunotherapy, coupled with log-ratio transformations to quantify ecosystem-level restructuring of the gut microbiota [Morton et al., 2019]. This methodological decision was driven by the compositional nature of microbiome data, where log-ratios mitigate feature interdependencies while preserving ecological interpretability, and their inherent robustness to technical biases such as uneven Gram-negative/positive bias during DNA extraction or sequencing depth. This framework aligns with contemporary paradigms in microbiome research, where holistic community dynamics supersede isolated taxonomic signatures [Gloor et al., 2017].

The identification of OGUs associated with immunotherapy outcomes followed an established analytical framework from our previous works [Olekhnovich et al., 2023; Zakharevich et al., 2024]. Differential rankings were first performed using Songbird v1.0.3 [Morton et al., 2019] to identify OGUs showing relative abundance variations between experimental groups, applying a conservative absolute differential value threshold of > 0.3. For candidate OGUs that met this criterion, we subsequently calculated log-ratio abundances according to the principle described in the works of Morton et al. (2019) and Fedarko et al. (2020). The statistical significance of the division of patients by response based on log ratios was determined through Wilcoxon rank-sum tests with alpha threshold = 0.05 implemented in the R statistical environment v4.4.2. The biomarker selection process incorporated stringent cross-validation criteria to ensure robust identification of microbial signatures. OGUs demonstrating consistent positive associations with therapeutic response across multiple datasets were retained as potential beneficial biomarkers, while any evidence of negative association with treatment outcome in any dataset resulted in automatic exclusion regardless of other positive associations. This approach enabled simultaneous identification of two clinically meaningful biomarker categories: microbial taxa positively correlated with successful immunotherapy outcomes and those associated with adverse therapeutic responses. The methodology emphasizes reproducibility through multi-dataset validation and maintains rigorous standards for biomarker qualification by requiring consistent directional effects across independent cohorts. Correlation analysis between regression coefficients and OGU prevalence was performed similarly to the analysis with MaAsLin2 regression coefficients. Additional validation of the relationship was performed through the glm package using the ranova function from the lmer package to test the influence of the dataset variable.

### Validation of OGU as a candidate marker for stratifying patients by response to immunotherapy

Determined R/NR biomarkers were used to calculate log ratios, followed by statistical assessment using the lmer function from the lmerTest package v3.1.3 [Kuznetsova et al., 2017] in the R statistical environment (log_ratio ∼ response + (1 | dataset)). Model residuals were inspected for normality through visual examination. Bootstrap validation was performed using the lmeresampler package. We applied a case bootstrap approach with 1000 iterations, resampling both individual observations and groups, to generate 95% percentile confidence intervals for the fixed effects estimates. Additional testing of marker OGU included investigating the ability of calculated log ratios to separate experimental groups using logistic regression using the method of “leave-one group out” (LOGO) cross validation using the tidymodels package v1.3.0. Rahman et al. (2023) demonstrated the effectiveness of using log-ratios based log regression in machine learning models.

### OGUs origin prediction

The presence of non-gut microbes in the stool microbiome, such as those frequently detected on the oral means, has been associated with a degradation of the gut microbiota [Liao et al., 2024]. This phenomenon frequently results in adverse outcomes, including various illnesses. The identification of non-gut microbes was conducted using published microbial metagenome-assembled genomes (MAGs) from body sites (oral, skin, vagina) [Pasolli et al., 2019] and food [Carlino et al., 2024]. The body site MAGs were retrieved from http://segatalab.cibio.unitn.it/data/Pasolli_et_al.html. A total of 9,412 genomes were selected based on specific criteria, including those from the “Body Site” category, such as the airways, nasal cavity, oral cavity, skin, and vagina. Genomes categorized in the “Age category” as “school age” (12-19 years), “adult” (19-65 years) and “senior” (65-70 years) were included in the analysis. Genomes from newborns (<1 year) or children (1-12 years) were excluded, as the gut microbiota typically reaches an adult-like phylogenetic composition by 3 years of age [Yatsunenko et al., 2012]. The selected genomes were then clustered with our reconstructed catalog via dRep v3.4.5 [Olm et al., 2017] with 95% nucleotide identity and -ignoreGenomeQuality parameter. Genomes that passed the similarity cutoff to body site MAGs were labeled from the same source. The food MAGs were obtained from https://zenodo.org/records/13285428, where the cFMD_mags.tar.gz archive contains the folder cFMD_mags_prok with prokaryotic genomes. These 10,112 MAGs were pre-clustered among themselves using dRep v3.4.5 with 95% nucleotide identity and -ignoreGenomeQuality parameter, yielding a total of 983 genomes. The subsequent source identification process was consistent with that employed for body MAGs.

### Association analysis of OGUs R/NR marker sets

GSEA was used as a summarizing tool to study the statistical relationships of OGU marker sets with immunotherapy response. This analysis was implemented using the GSEA function from the clusterProfiler package (v4.16.0) [Xu et al., 2024] and examined relationships across taxonomy, gene sets, and other traits such as presence in food or body site. For this analysis, items for each trait were ranked based on the p-value multiplied by the sign of the Songbird regression coefficient (R/NR + k), where a negative sign indicates association with the NR group and a positive sign indicates association with the R group. GSEA analysis results were visualized using the GseaVis package v0.1.1 [Zhang et al., 2025].

### Functional annotation of OGUs marker sets

Functional annotation of the OGU markers was performed using MetaCerberus v1.4.0 [Figueroa et al., 2024] against the CAZy [Drula et al., 2022] and KEGG [Kanehisa et al., 2025] databases. To identify significant functional differences between marker sets, we employed a multi-tiered statistical approach beginning with Wilcoxon rank-sum tests to compare gene abundance distributions. Subsequently, Fisher’s exact tests with Benjamini-Hochberg FDR correction < 0.05 were applied to identify significantly enriched functional features between experimental groups. Finally, GSEA was implemented to evaluate coordinated changes at the level of CAZy enzyme classes and KEGG pathways.

## Declarations

### Ethics approval and consent to participate

For the analysis, sample identifiers from the SRA archive and publicly accessible metadata from published studies were employed.

### Consent for publication

The data used in this study were sourced from SRA/NCBI. All original studies obtained informed consent from participants for the publication of anonymized data. Our analysis did not require additional ethical approval, as only de-identified datasets were utilized from open sources.

### Data availability

In this study, we used open access data from the NCBI/EBI Sequence Read Archives, identified by the following BioProjects accession numbers: PRJNA397906, PRJEB22893, PRJNA399742, PRJNA678737, PRJNA672867, PRJNA770295, PRJEB43119, PRJNA762360, PRJNA1011235, PRJNA928744, PRJNA615114, PRJNA866654, PRJNA494824, PRJEB49516. The pipeline for OGU catalog assembly and the sampleid.txt file containing the SRA/NCBI accession numbers used are available at: https://github.com/JeniaOle13/Cancer_MAGs. All initial MAGs sequences were deposited in NCBI under accession PRJNA1196825. Source code and Quarto report available at https://github.com/JeniaOle13/cancer-biomarkers. A preprint of this work is available on the bioRxiv https://www.biorxiv.org/content/10.1101/2025.05.07.652660.abstract.

### Authors’ contributions

VAO performed the computational analysis, including metagenomic data processing, microbial feature identification, and taxonomic profiling. VAO performed cross-database comparisons to trace microbial origins and contributed to the writing and editing of the manuscript. EIO conceived and designed the study, secured funding, and supervised all aspects of the research. EIO performed the statistical modeling and biomarker validation framework, led the biological interpretation of the results, and wrote and revised the manuscript, approving the final version.

## Supplementary information

**Table S1.** Genome quality statistics for the non-redundant catalog of 3,816 MAGs, including completeness, contamination, and taxonomic classification.

**Table S2.** Clinical and technical metadata for the curated cohort of 624 baseline stool metagenomes from 11 studies, including patient response status, cancer type, age, gender, and dataset source.

**Table S3.** Normalized relative abundance matrix of 3,816 OGUs across 624 samples, generated by mapping reads to the custom MAG catalog for consistent taxonomic profiling. Table S4. Differential abundance analysis with MaAsLin2 of microbial taxa associated with immunotherapy response.

**Table S5.** Dataset-specific results from the correlation analysis between microbial prevalence and Songbird-derived association coefficients with immunotherapy response.

**Table S6.** Complete set of 350 marker OGUs identified by Songbird differential ranking, annotated with their direction of association (R/NR) and regression coefficients.

**Table S7.** Sample-level log-ratio values derived from the 350 marker OGUs, serving as the core predictive feature for immunotherapy response stratification.

**Table S8.** Results of the exogenous origin analysis showing catalog OGUs that cluster at 95% Average Nucleotide Identity (ANI) with reference MAGs from oral, other body sites, and food sources.

**Table S9.** Differential enrichment of CAZy families in the genomes of R- versus NR-associated marker OGUs (Fisher’s exact test, FDR-corrected).

**Table S10.** GSEA results showing GH families coordinately enriched in the R-associated marker OGUs.

**Table S11.** Differential abundance of KO groups in the genomes of R- versus NR-associated marker OGUs (Fisher’s exact test, FDR-corrected).

**Table S12.** Pathway-level GSEA of KEGG metabolic pathways in the genomes of R- versus NR-associated marker OGUs.

**Figure S1.** Validation of the log-ratio biomarker for immunotherapy response. (A) Distribution of log-ratio values, calculated from marker OGUs, for R and NR patients. (B) LOGO cross-validation results assessing the predictive performance of the log-ratio-based biomarker. The line indicates the mean AUC across all datasets (0.67 ± 0.13).

**Figure S2.** Comprehensive performance metrics of the log-ratio biomarker across independent cohorts. The figure presents detailed prediction metrics from the LOGO cross-validation analysis. Metrics include the area under the receiver operating characteristic curve (AUC), accuracy, sensitivity, specificity, precision, and F1-score.

