## Supplementary figures and images for "Ecological and functional stratification of the stool microbiome predicts response to immune checkpoint inhibitors across cancer types"

### Supplementary Figure S1

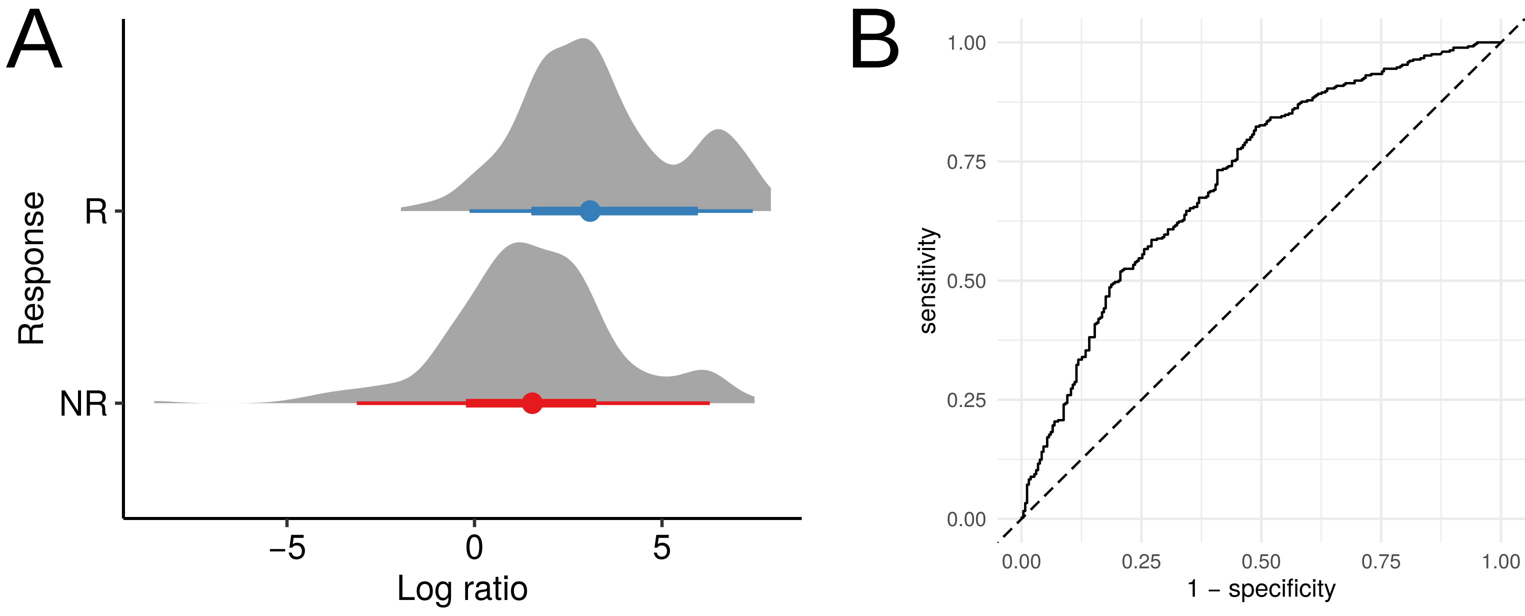

### Supplementary Figure S2

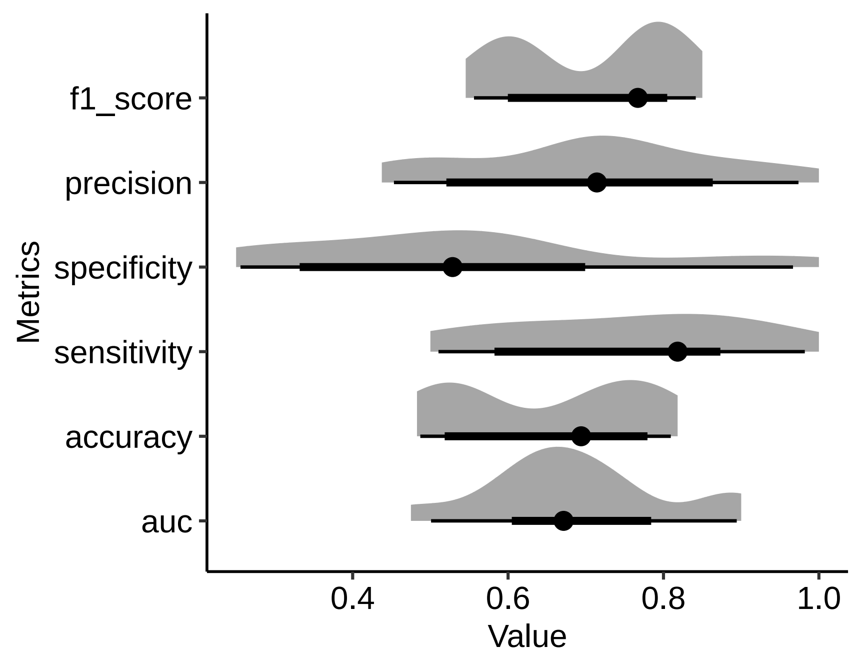
